# Pre-Innervated Tissue Engineered Muscle Promotes a Pro-Regenerative Microenvironment Following Volumetric Muscle Loss

**DOI:** 10.1101/840124

**Authors:** Suradip Das, Kevin D. Browne, Franco A. Laimo, Joseph C. Maggiore, Halimulati Kaisaier, Carlos A. Aguilar, Zarina S. Ali, Foteini Mourkioti, D. Kacy Cullen

**Affiliations:** Center for Brain Injury & Repair, Department of Neurosurgery, Perelman School of Medicine, University of Pennsylvania, Philadelphia, PA; Center for Neurotrauma, Neurodegeneration & Restoration, Corporal Michael J. Crescenz Veterans Affairs Medical Center, Philadelphia, PA; Department of Bioengineering, School of Engineering and Applied Science, University of Pennsylvania, Philadelphia, PA; Department of Biomedical Engineering, University of Michigan, Ann Arbor, MI; Penn Center for Musculoskeletal Disorders, Department of Orthopedic Surgery, Perelman School of Medicine, University of Pennsylvania, Philadelphia, PA; Department of Cell and Developmental Biology, Perelman School of Medicine, University of Pennsylvania, Philadelphia, PA; Penn Institute for Regenerative Medicine, Musculoskeletal Program, Perelman School of Medicine, University of Pennsylvania, PA

**Author notes:** Corresponding author: D. Kacy Cullen, Ph.D. 105E Hayden Hall/3320 Smith Walk Dept. of Neurosurgery University of Pennsylvania Philadelphia, PA 19104.

## Abstract

Volumetric Muscle Loss (VML) is defined as traumatic or surgical loss of skeletal muscle tissue beyond the inherent regenerative capacity of the body, generally leading to a severe functional deficit. Autologous muscle grafts remain the prevalent method of treatment whereas recent muscle repair techniques using biomaterials and tissue engineering are still at a nascent stage and have multiple challenges to address to ensure functional recovery of the injured muscle. Indeed, appropriate somato-motor innervations remain one of the biggest challenges for both autologous muscle grafts as well as tissue engineered muscle constructs. We aim to address this challenge by developing Pre-Innervated Tissue Engineered Muscle comprised of long aligned networks of spinal motor neurons and skeletal myocytes. Here, we developed methodology to biofabricate long fibrils of pre-innervated tissue engineered muscle using a co-culture of myocytes and motor neurons on aligned nanofibrous scaffolds. Motor neurons lead to enhanced differentiation and maturation of skeletal myocytes *in vitro*. These pre-innervated tissue engineered muscle constructs when implanted *in vivo* in a rat VML model significantly increase satellite cell migration, micro-vessel formation, and neuromuscular junction density in the host muscle near the injury area at an acute time point as compared to non-pre-innervated myocyte constructs and nanofiber scaffolds alone. These pro-regenerative effects can potentially lead to enhanced functional neuromuscular regeneration following VML, thereby improving the levels of functional recovery following these devastating injuries.

## Introduction

Innervation plays a crucial role in the development, maturation and functioning of different muscles in the body and yet remains largely unexplored in tissue engineering studies related to cardiac, smooth or skeletal muscle. There are limited reports showing the importance of re-innervation in a transplanted heart or in a tissue engineered urinary bladder in order to achieve complete functional recovery^1–3^. Tissue engineered constructs typically lack preformed neural networks and depend on host-induced innervation to integrate with the native nerve supply. Although the concept of “pre-innervation” (akin to pre-vascularization), wherein appropriate neuronal populations are cultured along with relevant muscle cells has been established for a long time in vitro ^4–7^, it is yet to be studied in vivo in models of muscle injury. The present study is focused towards exploring the effect of pre-innervation in skeletal muscle tissue engineering using a severe muscle trauma model like Volumetric Muscle Loss (VML).

VML is defined as traumatic or surgical loss of a large mass of skeletal muscle tissue beyond the inherent regenerative capacity of the body, generally leading to a severe functional deficit^8^. Due to a significant loss of nerve and vascular supply accompanied by inflammation driven fibrosis the self-repair process is not adequate to generate sufficient new muscle in time to prevent a chronic scar^9,10^. Although VML is widespread among the civilian population, military personnel in particular are more prone to such damage due to combat related musculoskeletal injuries^11^. According to a recent study, 65% of soldiers who retired due to various injuries reported a muscle condition, while 92% of these cases included VML^12^.

Free functional muscle transfer (FFMT) is the preferred procedure to treat VML, which entails transfer of donor muscle along with nerve and blood vessels from another part of the body to the injury site to facilitate re-innervation and re-vascularization of the graft region^9,11,13^. Although FFMT remains the gold standard, its success is limited by donor site morbidity, long operative time and prolonged de-innervation of motor end plates in the donor muscle^14^. As an alternative approach, tissue engineered skeletal muscle constructs have been fabricated using scaffold based as well as scaffold-less technologies. Decellularized extracellular matrix (dECM) remain the most prevalent scaffold material used for muscle reconstruction following VML injuries^15,16^. Although dECM-based scaffolds recapitulate the native ECM composition, their efficacy is challenged by their fast resorption rates in the body and failure to provide topographical guidance to regenerating myofibers thereby leading to random fiber recruitment and subsequent scar tissue formation^17,18^. The ECM secreted by skeletal myofibers is arranged in a network of micro-nano fibrils which are aligned along the myofibers. Aligned nanofiber-based scaffolds accurately replicate the topographical cues of native ECM architecture and has been reported to promote aligned myogenesis, cell migration, survival and angiogenesis^19^.

Persistent inflammation, fibrosis, revascularization and reinnervation remain the major impediments to complete recovery of contractile function following major skeletal muscle trauma like VML^20–24^. Hence, in addition to topographical guidance, tissue engineered scaffolds must be myo-conductive, promote angiogenesis through either pre-vascularization or incorporation of vasculogenic accelerant, have immunomodulatory effect as well as facilitate reinnervation to overcome the pathophysiology of VML and achieve functional restoration. Direct administration of anti-inflammatory agents can reduce fibrosis but has been shown to hinder muscle regeneration^25,26^. Acellular scaffolds cannot address such multidimensional challenges and reinforces the necessity of incorporating multiple cell types in a scaffold^18^. Pre-vascularized constructs fabricated by coculture of myoblasts and endothelial cells on aligned nanofibrous scaffolds have been found to promote organized myofiber regeneration and vascular integration in murine VML model^27^. Although angiogenic and immunomodulatory strategies have been explored in VML repair, lack of innervation in an engineered muscle remains one of the major impediments to its success as a functional muscle replacement^28^. In the absence of neural innervation, the functional maturation of myofibers cannot proceed^29^. Hence it is imperative that tissue engineering strategies for muscle replacement should consider innervation as an essential component of the biofabrication process itself to facilitate maturation of myofibers in vitro as well as achieve robust host innervation in vivo following implantation in severe musculoskeletal injuries.

In the present study, we report the development of a novel pre-innervated tissue engineered muscle construct for application in VML repair (Fig 1). Aligned nanofiber sheets were used to coculture skeletal myocytes and spinal motor neurons and explore the effect of innervation on maturation of myocytes in vitro (Fig 1a). The bioscaffolds were further implanted in athymic rat model of VML and evaluated for cell survival, satellite cell proliferation, microvasculature and neuromuscular junction density (NMJs) at an acute time point (Fig 1b). This is the first report studying the acute effect of pre-innervated constructs on the regenerative micro-environment of injured muscle following severe muscle trauma.

**Fig. 1:**
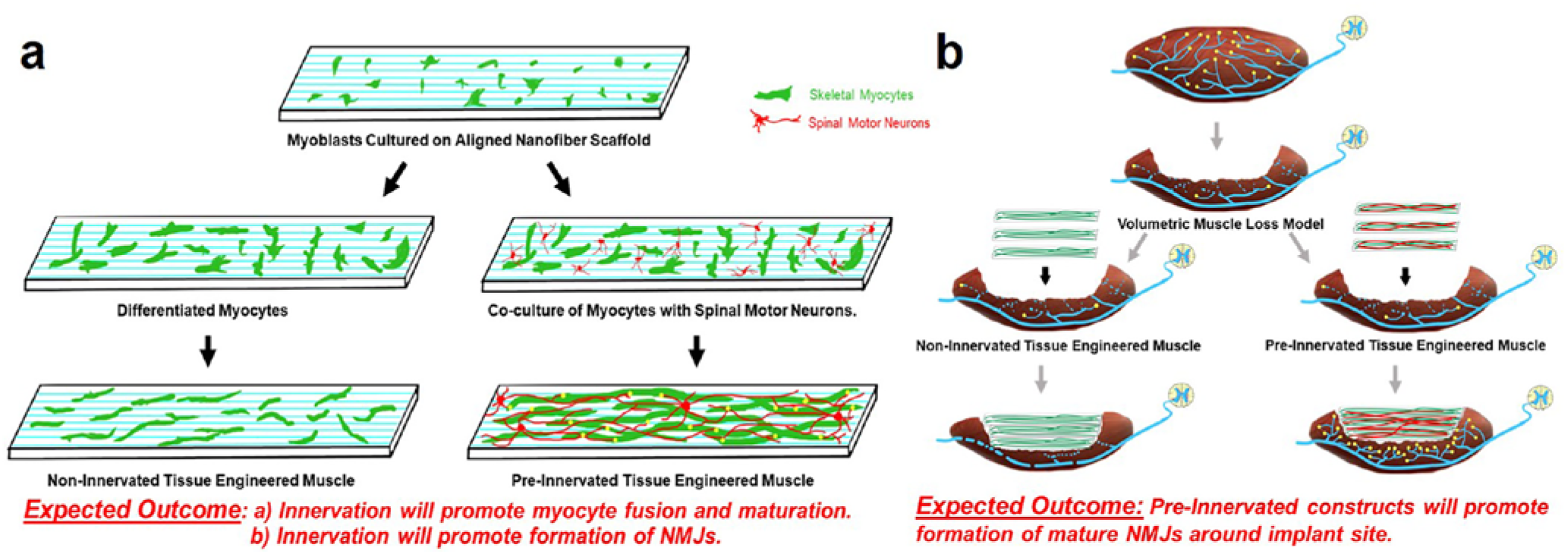
Concept of Pre-Innervated Tissue Engineered Muscle. The present study was focused on exploring the role of pre-innervation on myocytes in vitro and host neuromuscular environment in vivo following implantation. **a)** For in vitro studies, our overarching hypothesis were that innervation would augment skeletal myocyte fusion, maturation and formation of Neuromuscular Junctions (NMJs). **b)** Volumetric Muscle Loss (VML) is defined as frank loss of muscle volume that is accompanied by chronic motor axotomy leading to denervation of the injured area. We used a standardized rat model of VML where >20% of the Tibialis Anterior (TA) muscle volume was resected to create a defect leading to potential damage to intramuscular branches of the host nerve and loss of motor end plates (or NMJs) near the injury area. For in vivo studies, our overarching hypothesis were that implantation of pre-innervated constructs would enhance Acetylcholine Receptor (AchR) clustering and promote innervation of AchRs (mature NMJs) near the implant site at acute time point.

## Results

### Pre-Innervation promotes myocyte fusion and formation of NMJs in vitro

Mouse skeletal myoblast cell line C2C12 were cultured on aligned polycaprolactone (PCL) nanofiber scaffolds and allowed to differentiate. Differentiated myofibers were found to align along the direction of nanofibers as observed by staining for F-actin (Fig 2a). Similarly, spinal motor neurons cultured on the nanofibers exhibited axons aligning along the nanofiber orientation (Fig 2b). Subsequently, both motor neurons and myocytes were cocultured on the nanofiber scaffolds. Motor neuron-myocyte coculture led to formation of thick intertwined nerve-muscle bundles aligned along the nanofibers (Fig 2c-c″). Within 7 days of coculture on the nanofiber scaffolds, NMJs were observed by colabelling for presynaptic marker Synaptophysin and Bungarotoxin mediated identification of post synaptic Acetylcholine Receptors (AchR) (Fig 3a-a′). Motor neurons were also found to promote myocyte maturation and fusion in vitro leading to significantly higher myocyte fusion index (MFI) as compared to myocyte only cultures (Fig 3b-c). Taken together, these data clearly demonstrates that innervation not only leads to NMJs in vitro but also facilitates myocyte maturation.

**Fig. 2:**
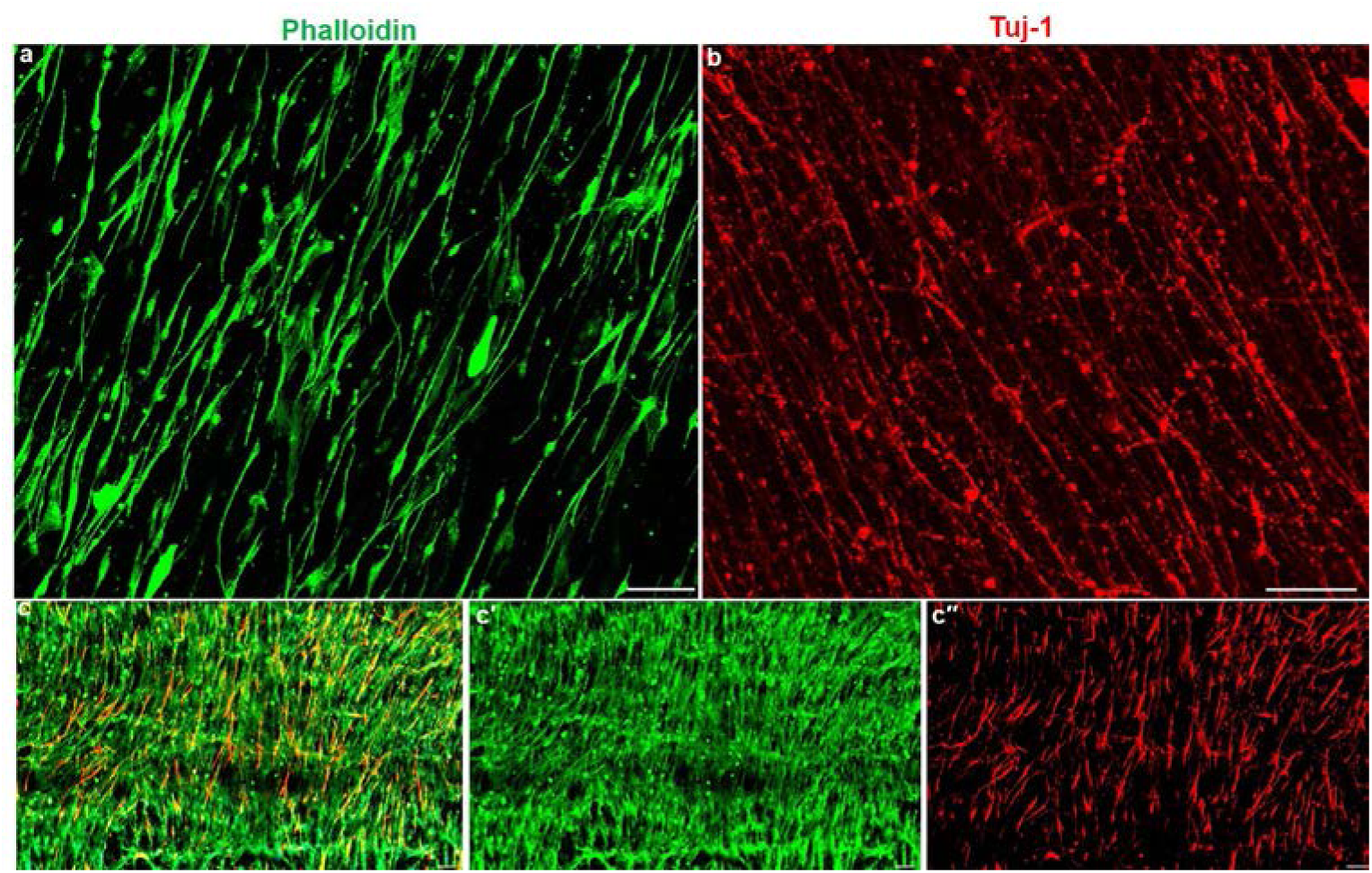
Coculture of Motor Neuron-Myocyte on Aligned Nanofiber Scaffolds. **a)** Skeletal myoblast cell line C2C12 was cultured and allowed to differentiate on aligned nanofiber scaffolds. Mature myofibers were observed to align along the nanofibers when stained for F-Actin (Phalloidin-488). Scale bar **–**200µm. **b)** Spinal motor neurons cultured on the nanofiber scaffolds aligned along the direction of nanofibers as observed by staining with pan-axonal marker Tuj-1 (Red). Scale bar – 200µm. **c-c″)** Coculture of motor neurons and myocytes on the nanofiber scaffolds resulted in formation of aligned neuromuscular bundles as observed by labelling for **c′)** F-Actin (Phalloidin-488) and **c″)** Tuj-1 (Red). Scale bar – 100µm.

**Fig. 3:**
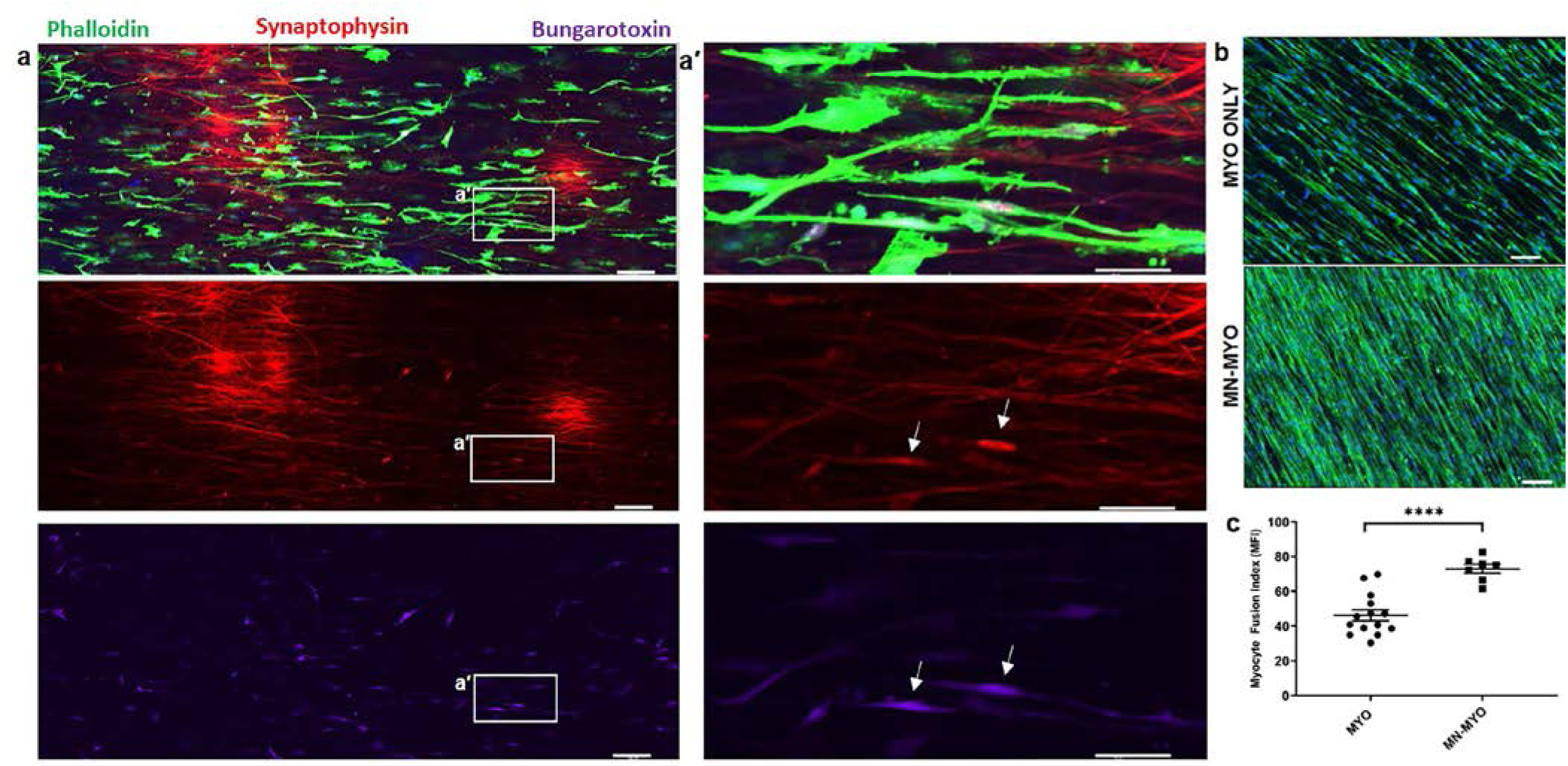
Innervation of Myocytes and Effect of Motor Neurons on Myocyte Maturation *In Vitro*. **a)** Rat spinal motor neurons were introduced on a bed of myofibers differentiated on aligned nanofiber sheet for 7 days and subsequently cocultured for another 7 days leading to innervation of the skeletal myofibers. Scale bar – 100µm. **a′)** Higher magnification view of the area marked by white box reveals structures colabelling for presynaptic marker (Synaptophysin) and Acetylcholine Receptor (AchR) clusters (Bungarotoxin) indicating formation of mature neuromuscular junctions *in vitro* (indicated by white arrows). Scale bar – 50µm. **b)** Myocytes exhibited greater fusion and bundling when cocultured with motor neurons (MN-MYO) as compared to monoculture (MYO). Scale bar – 100µm**. c)** Myocyte Fusion Index (MFI) was calculated from multiple cultures (n ≥ 6), and coculture with motor neurons was found to significantly enhance MFI. For indicated comparison: p ≤ 0.0001 (****). Error bars represent standard error of mean.

### Bioscaffold implantation in athymic rat model of VML

The tibialis anterior (TA) muscle of athymic rats was exposed and a 10 mm×7 mm×3 mm (length × width × depth) segment of the muscle was excised corresponding to ~20% of gross muscle weight to create a VML model (Fig 4a-b). The animals were randomized into the following repair groups: (I) three stacked nanofibrous sheets (per animal) containing coculture of motor neurons and myocytes (MN-MYO; n=6); (II) three stacked nanofibrous sheets containing myocytes only (MYO; n=4); (III) three stacked acellular nanofibrous sheets only (SHEETS; n=3) and (IV) NO REPAIR (n=5) (Fig 4c). At terminal time point of 7 days post implant, the nanofiber sheets were visible upon TA exposure and appeared intact (Fig 4d). Further, the graft area in the No Repair group appeared to be recessed and atrophied as compared to the Repair groups (Fig 4e-f).

**Fig. 4:**
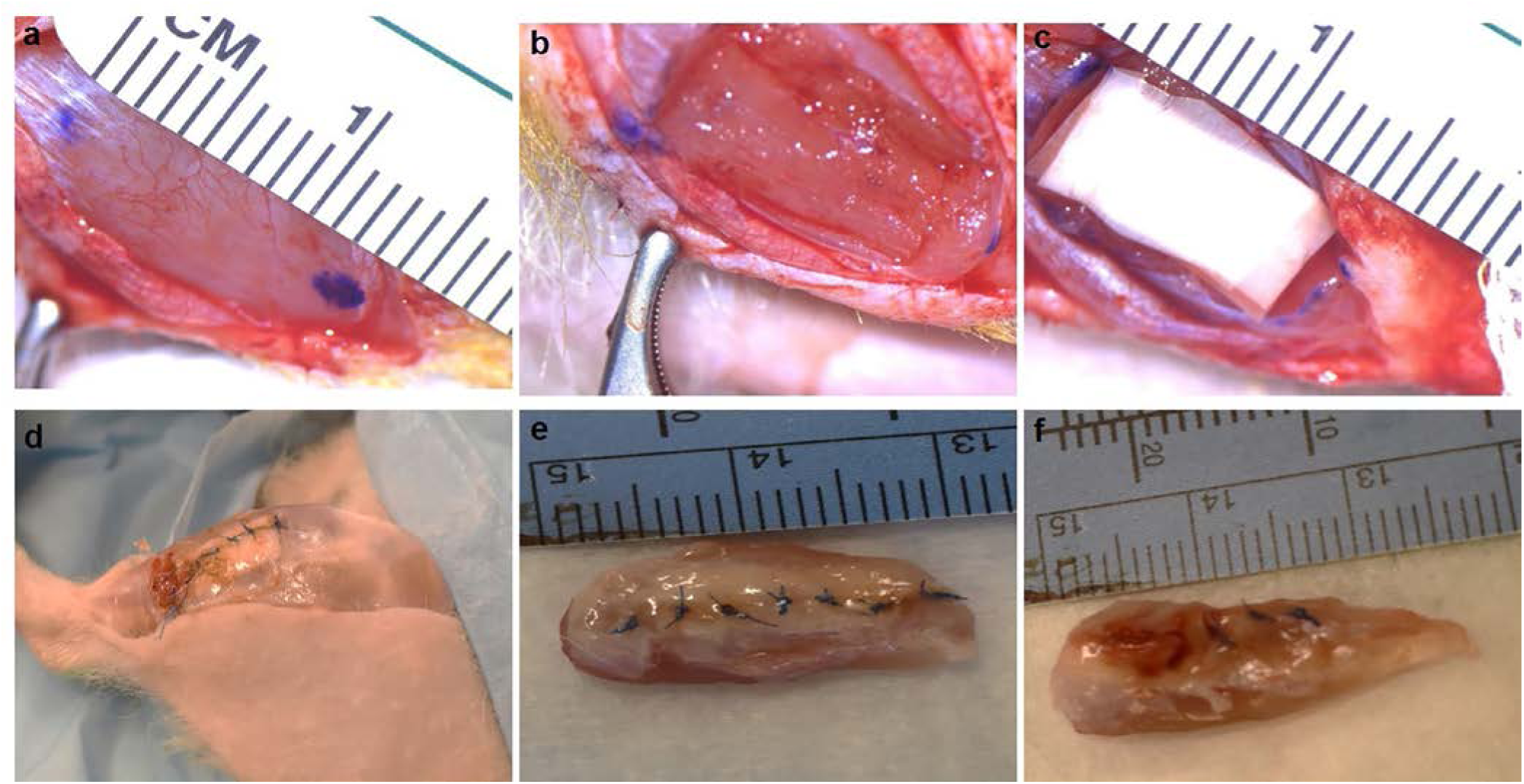
Bio-Scaffold Implantation in VML Model. **a-b**) Surgical resection of TA muscle to create VML model in rats. **c**) Implant of cell-laden nanofiber sheets in muscle defect. Scaffold and overlying fascia secured with sutures. **d-f**) At 7 days post-implant, animals were sacrificed, and TA muscle was excised. Nanofiber sheets were seen in Repair Group (**e**) whereas injury site was recessed in No Repair group (**f**).

### Evaluation of acute cell survival in bioscaffolds upon implantation in VML model

All animals were sacrificed after 7 days and the whole anterior muscle compartment of the hind limbs were fixed in paraformaldehyde. The muscles were cryopreserved, embedded in OCT, sectioned and stained. Immunohistochemical analysis of cross sections of the injury/repair sites were performed to detect implanted cells on the nanofiber sheets and assess overall muscle health. We found that the nanofiber sheets were intact after 7 days in all the animals. Myocytes positive for Phalloidin (F-actin) were observed in both MN-MYO and MYO groups (Fig. 5a-b). Animals implanted with SHEETS only did not show significant Phalloidin+ cells within the implant region while the NO REPAIR group was left with a gap that was eventually found to be filled with infiltrating cells (Fig. 5c-d). Interestingly, axons positive for motor neuron marker Choline Acetyl Transferase (ChAT) and Neurofilament (NF-200) were observed within the nanofiber sheets in animals implanted with neuron-myocyte cocultures (MN-MYO) (Fig. 5a) whereas no axons/neurons were found within the injury/repair site in other groups. Longitudinal sections from the MN-MYO group revealed thick elongated myocytes and motor axons within the implanted nanofiber sheets (Fig. 6). These results indicate the survival of implanted motor neurons and myocytes at acute time point following a VML repair.

**Fig. 5:**
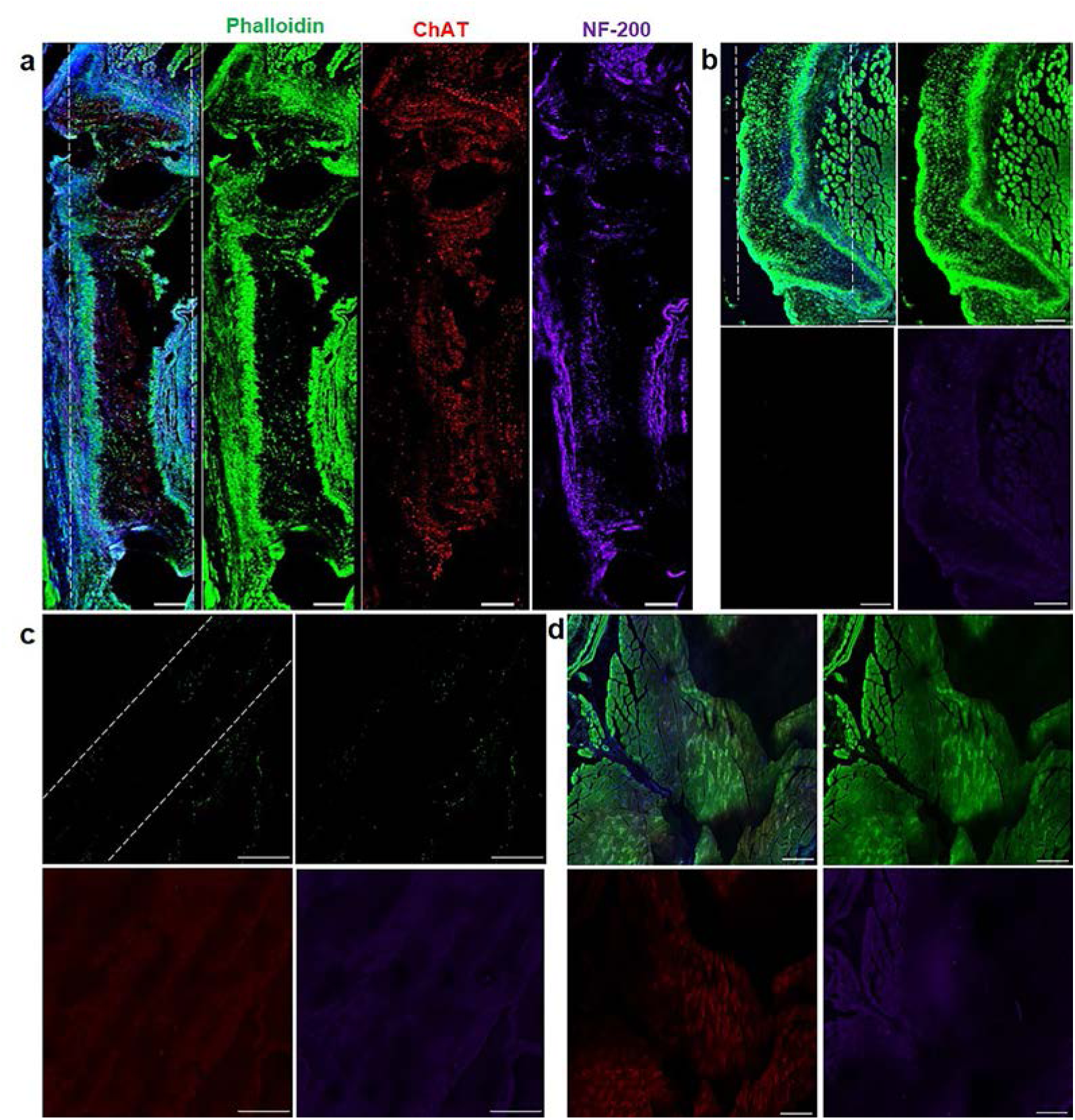
Acute Survival of Implanted Cells *In Vivo*. **a-d**) Cross section images of the repair/injury site from animals implanted with nanofibers comprised of **a**) motor neurons + myocytes (MN-MYO), **b**) myocytes only (MYO), **c**) acellular Sheets and **d**) No Repair. The sections showed presence of axons (ChAT+ & NF-200+) on the nanofiber sheets in the MN-MYO group whereas the other groups were negative for axonal markers at the injury site. The nanofiber sheets on both the MN-MYO and MYO groups showed presence of Phalloidin+ myocytes. Dashed lines indicate margins of nanofiber sheets. Scale bar **–**200µm.

**Fig. 6:**
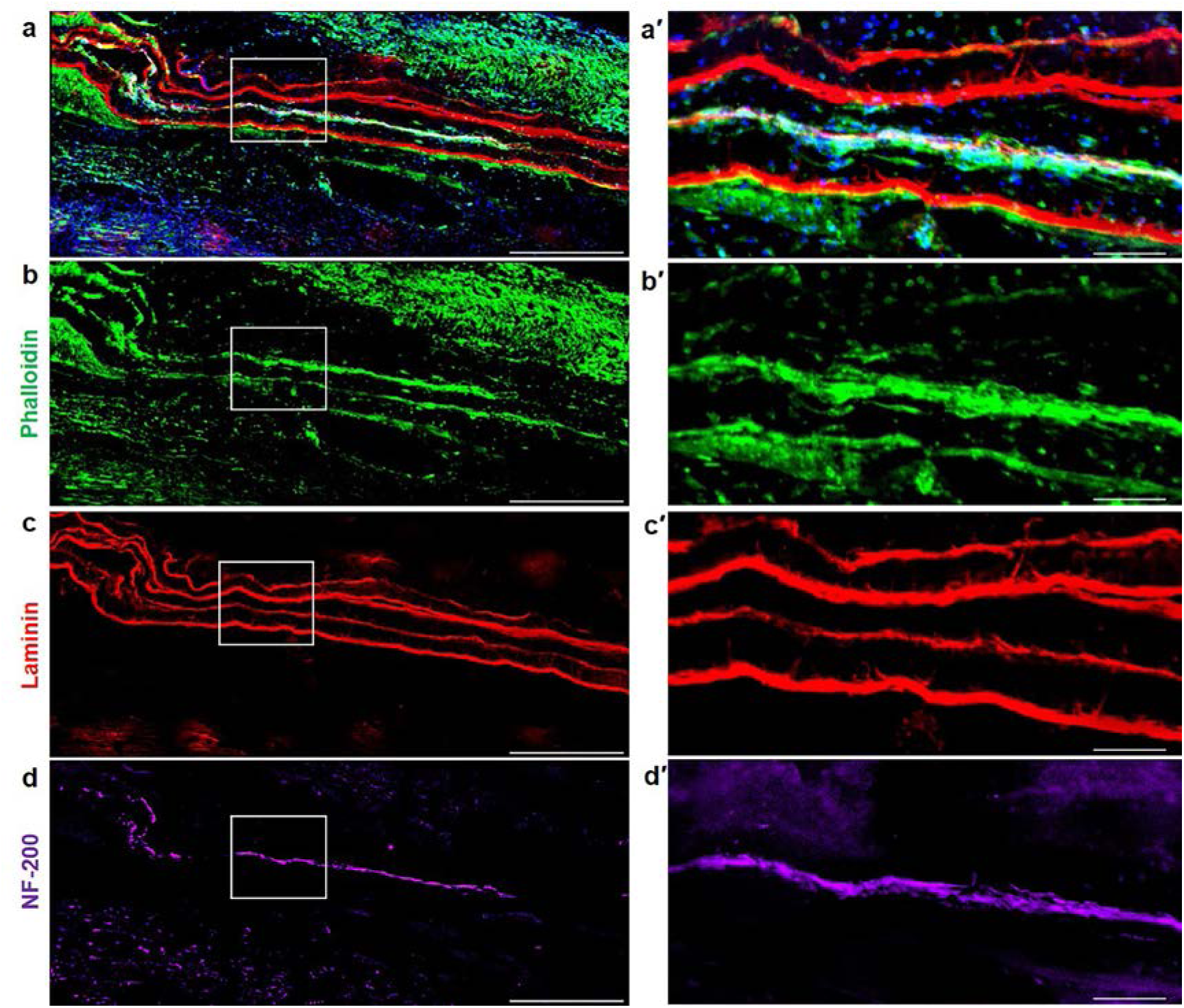
Cellular and Morphological Evaluation of Pre-Innervated Constructs at Acute Time Point Following Implantation in a VML Model. **a-d**) Longitudinal sections near the repair site of animals implanted with nanofibers with motor neurons + myocytes (MN-MYO). The nanofiber sheets were coated with Laminin prior to culturing cells and hence the stacked sheets were identified based on Laminin stain (red). Scale: 500µm. **a′-d′)** Magnified view of the region inside the white box. Thick bundles of myocytes (Phalloidin: green) and motor axons (NF-200: purple) were observed within the stacked nanofiber sheets. Scale: 100µm.

### Pre-Innervated constructs promote satellite cell migration near injury area

Satellite cells are resident myogenic precursor cells essential for muscle regeneration^30^. Activation and mobilization of satellite cells to the sites of injury is a major contributor to the regenerative capability of skeletal muscle^31^. Satellite cell migration near the injury area was observed across all groups by staining with Pax7 (Fig 7a-d). Pax7^+^ nuclei located on the periphery of Skeletal Muscle Actin^+^ myofibers and colabelling with pan-nuclear marker DAPI were identified as satellite cells (Fig 7e). Importantly, the pre-innervated MN-MYO group exhibited significantly higher satellite cell proliferation near the injury area as compared to other groups confirming that innervated constructs can trigger satellite cell migration and potentially facilitate muscle regeneration.

**Fig. 7:**
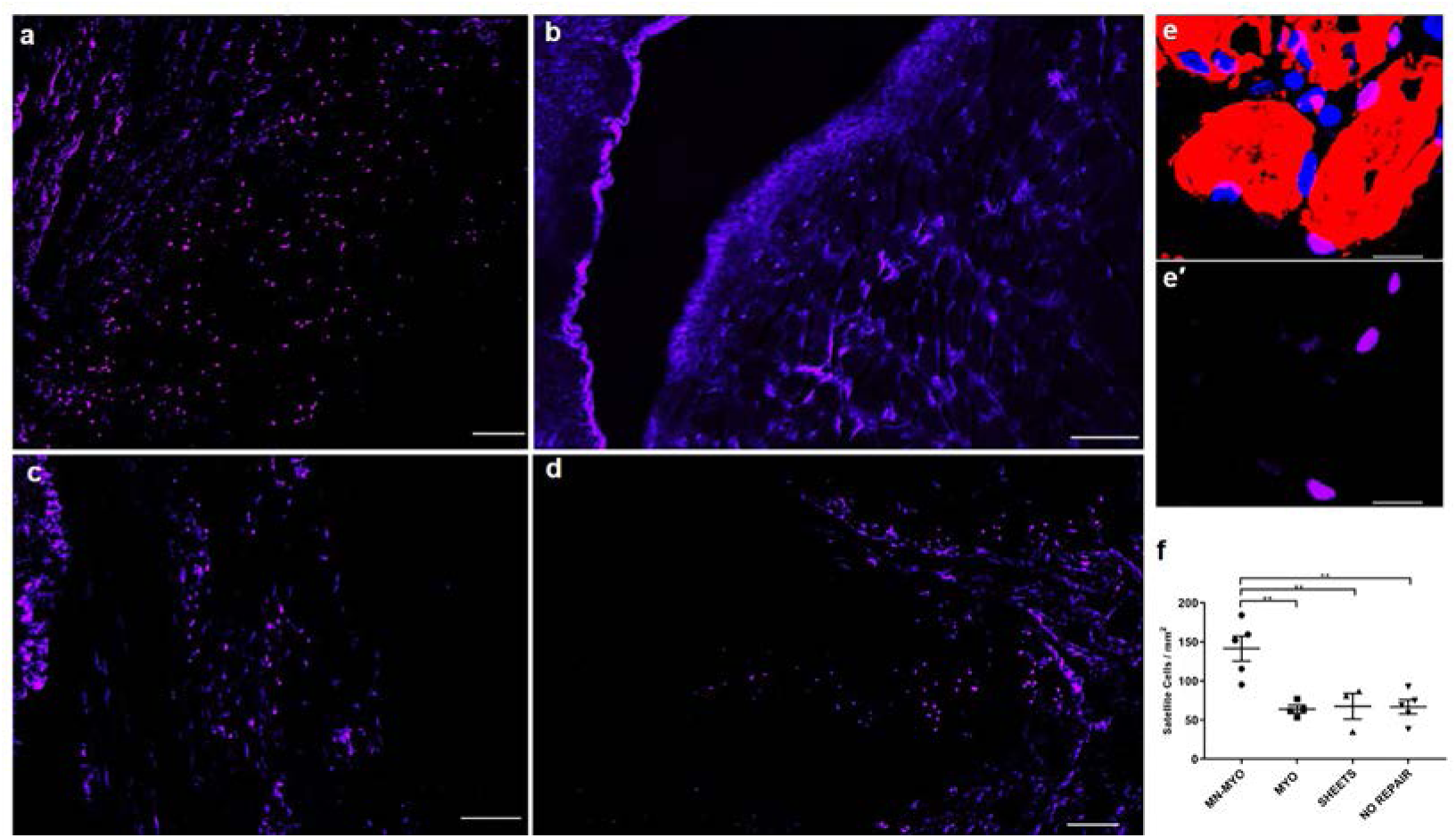
Satellite cell migration near injury area following VML. a-d) Muscle satellite cells near the injury area were identified by staining for satellite cell marker – Pax 7 (Purple) in a) MN-MYO; b) MYO; c) Sheets; d) No repair groups. Scale bar – 100µm. e-e′) Representative image of a higher magnification view of satellite cells. Pax 7+ nuclei (Purple) located on the periphery of Skeletal Muscle Actin+ (Red) myofiber and colabelling with pan-nuclear marker DAPI (Blue) were identified as satellite cells. Scale bar – 10µm. f) Satellite cell density near the injury area (5mm^2^) was counted across MN-MYO (n=5), MYO (n=4), Sheets (n=3) and No Repair (n=5) groups. Mean satellite cell density of each group were as follows: MN-MYO – 141.4; MYO – 64.06; Sheets – 67.75; No Repair – 66.85. For indicated comparisons the individual p-values were as follows: MN-MYO vs MYO – p=0.0029 (**); MN-MYO vs Sheets – p=0.0081 (**); MN-MYO vs No Repair – p= 0.0024 (**). Error bars represent standard error of mean.

### Pre-Innervated constructs lead to increased microvasculature near injury area

Revascularization of the injured area/implant is critical for survival of implanted cells and integration with host vascular system. Vascularization near the injury area was evaluated by staining tissue sections with endothelial cell specific marker-CD31 and Smooth Muscle Actin (SMA) (Fig 8). CD31^+^/SMA^+^ structures with a visible lumen and cross-sectional area greater than 50µm^2^ were defined as microvessels (Fig 8e). Although none of the groups exhibited migration of endothelial cells within the implanted sheets at this early time point (7 days), remarkably, the MN-MYO group showed presence of microvessel-like structures adjacent to the injury site while other groups had more punctate CD31^+^ cells (Fig. 8). The presence of significantly higher microvasculature around the injury area in MN-MYO group suggests that innervated tissue engineered muscle constructs can potentially augment revascularization following VML repair.

**Fig. 8:**
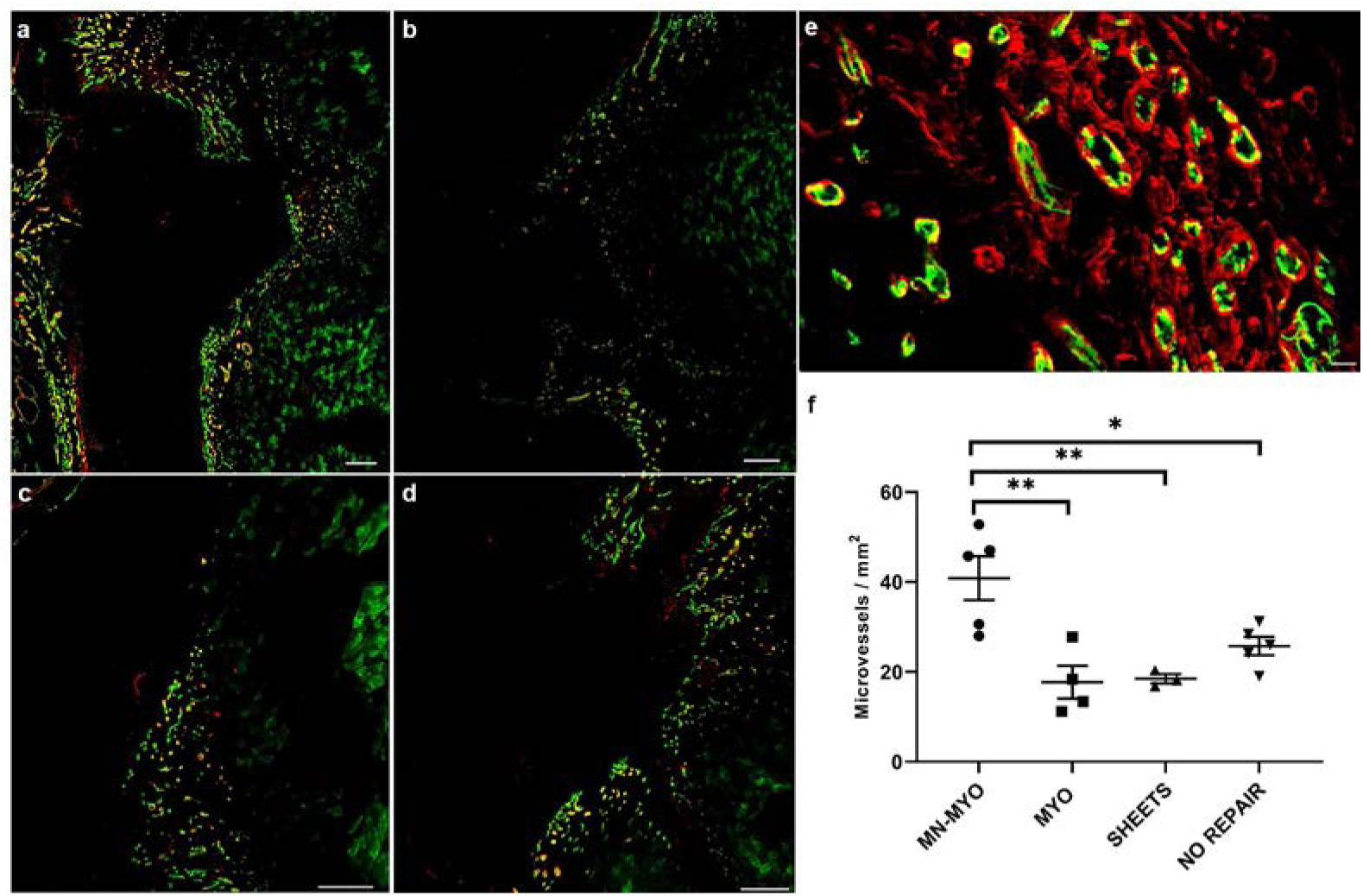
Micro-vessel density near injury area following VML. a-d) Endothelial cells and micro-vasculature near the injury area were identified by staining for endothelial cell marker – CD31 (Green) and Smooth Muscle Actin (Red) in a) MN-MYO; b) MYO; c) Sheets; d) No repair groups. Scale bar – 200µm. e) Representative image of a higher magnification view of endothelial cells and micro-vessels. Structures expressing CD31 (Green) and Smooth Muscle Actin (Red) with a visible lumen and an area > 50µm^2^ were defined as micro-vessels. Scale bar – 10µm. f) Micro-vessel density near the injury area (5mm^2^) was counted across MN-MYO (n=5), MYO (n=4), Sheets (n=3) and No Repair (n=5) groups. Mean microvessel density of each group were as follows: MN-MYO – 40.84; MYO – 17.7; Sheets – 18.47; No Repair – 25.76. For indicated comparisons the individual p-values were as follows: MN-MYO vs MYO – p=0.0024 (**); MN-MYO vs Sheets – p=0.0061 (**); MN-MYO vs No Repair – p= 0.0321 (*). Error bars represent standard error of mean.

### Enhanced Acetylcholine Receptor (AchR) expression following implantation of Pre-Innervated constructs

Acetylcholine Receptor (AchR) clusters have major implications in formation and maintenance of motor end plates during muscle development as well as regeneration^32,33^. Indeed, bungarotoxin staining, a known marker of nAchR a7 receptors^34^, showed the presence of pretzel-shaped AchR clusters around the injury area across all groups (Fig 9a-e). A count of AchR cluster near the injury area revealed that the MN-MYO group had significantly more AchR clusters than the other groups (Fig. 9f).

**Fig. 9:**
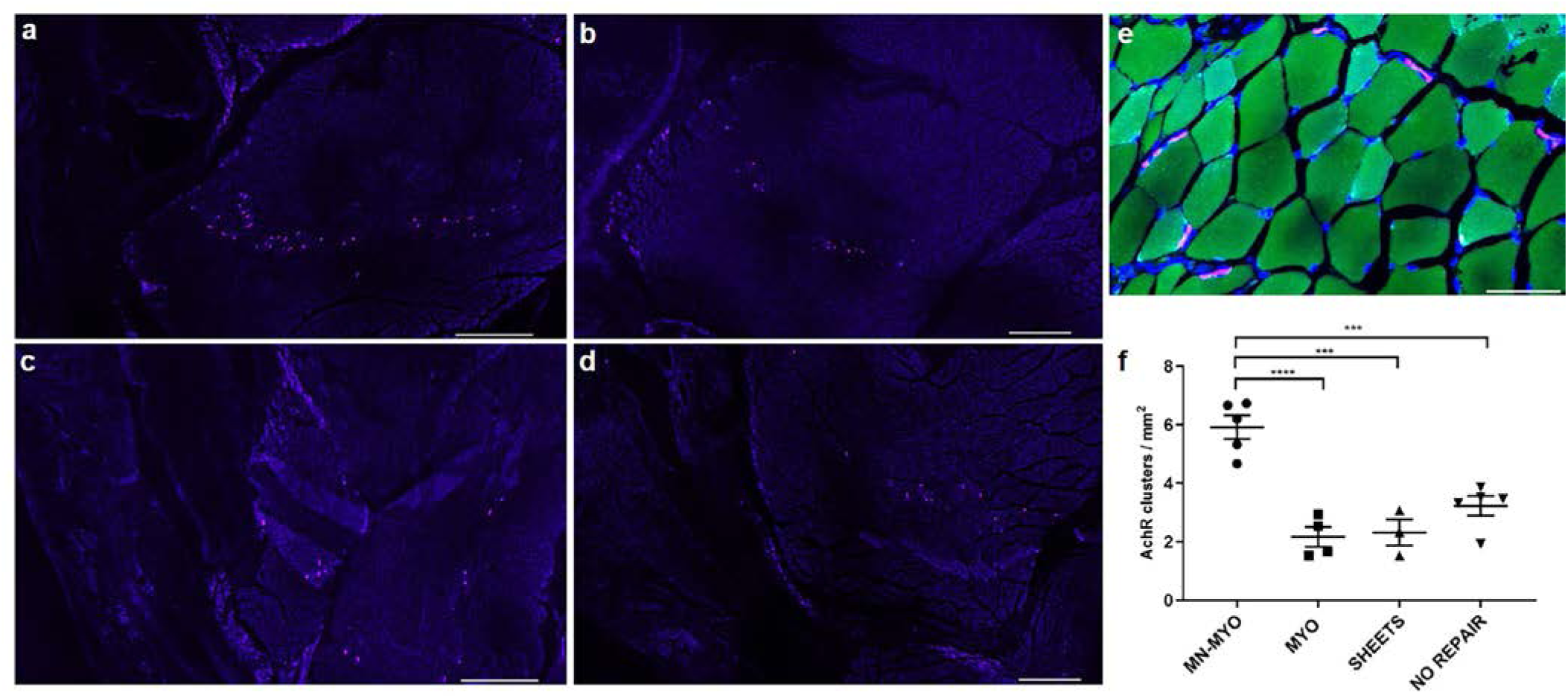
Acetylcholine Receptor (AchR) Clusters near injury area following VML. a-d) AchR clusters near the injury area were identified by staining with Bungarotoxin (Purple) in a) MN-MYO; b) MYO; c) Sheets; d) No repair groups. Scale bar – 500µm. e) Representative image of a higher magnification view of pretzel shaped AchR clusters (Purple) on the periphery of muscle fibers (Phalloidin-488). Scale bar – 50µm. f) AchR cluster density near the injury area (5mm^2^) was counted across MN-MYO (n=5), MYO (n=4), Sheets (n=3) and No Repair (n=5) groups. Mean AchR cluster density of each group were as follows: MN-MYO – 5.92; MYO – 2.167; Sheets – 2.311; No Repair – 3.227. For indicated comparisons the individual p-values were as follows: MN-MYO vs MYO – p<0.0001 (****); MN-MYO vs Sheets – p=0.0001 (***); MN-MYO vs No Repair – p= 0.0006 (***). Error bars represent standard error of mean.

### Pre-Innervated constructs promote formation of mature NMJs near injury area

Although AchR clustering is indicative of motor end plate health, they are not always the points of innervation or NMJs. Mature NMJs are indicative of muscle health and their loss has been implicated in neuromuscular degeneration associated with inflammation, denervation and atrophy^35^. Mature NMJs were identified as pretzel shaped structures which were colabelled with presynaptic marker Synaptophysin and postsynaptic AchR marker (Bungarotoxin) (Fig 10 a-e′). The percentage of AchR clusters near the injury area which were positive for Synaptophysin was calculated to quantify the amount of mature NMJs (Fig 10f). Pre-innervated constructs (MN-MYO) were found to have significantly higher percentage of mature NMJs as compared to other groups indicating the potential role of pre-innervation in augmenting formation of mature NMJs following implantation in VML model (Fig 10f).

**Fig. 10:**
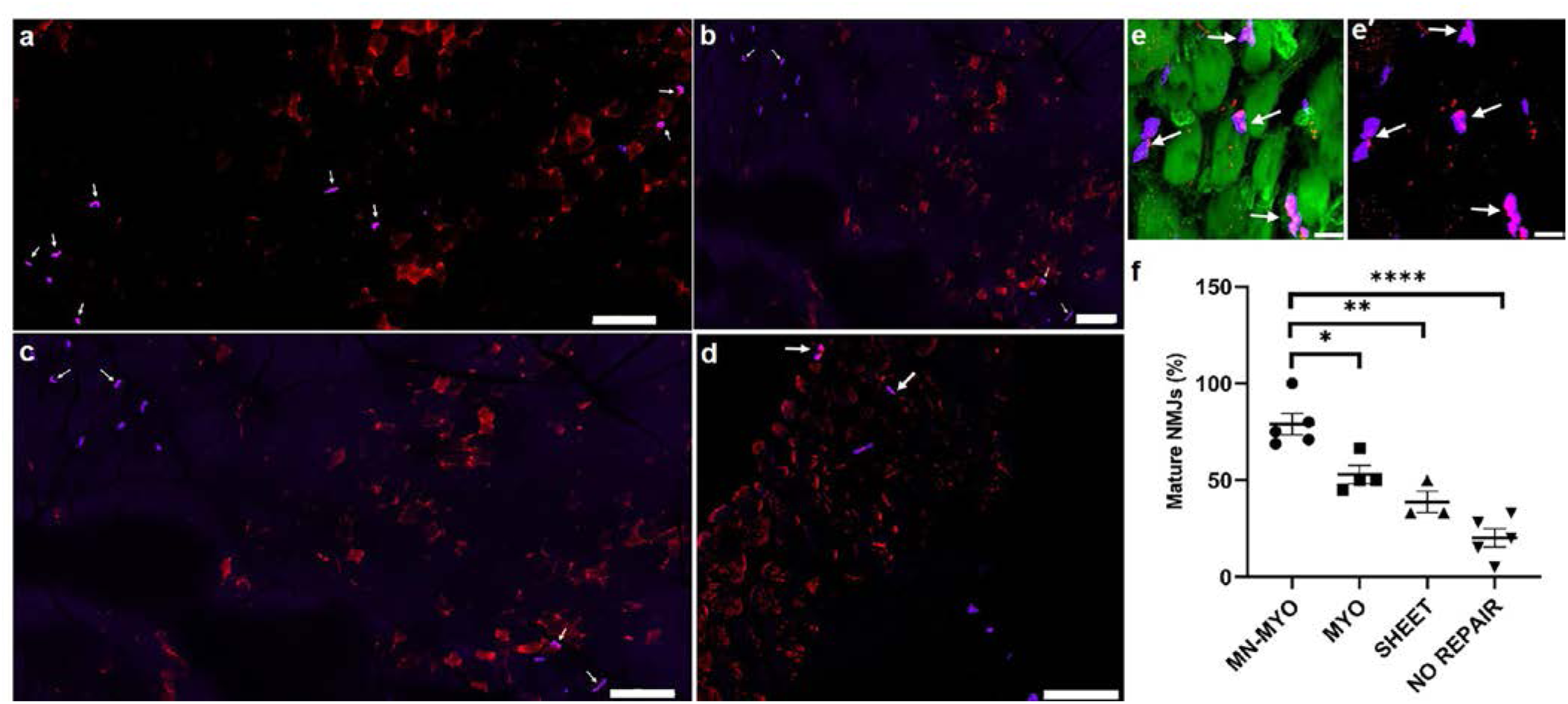
Pre-Innervation promotes mature NMJs formation near injury area following VML. a-d) Mature NMJs near the injury area were identified by double staining with Bungarotoxin (Purple) and presynaptic marker Synaptophysin (Red) in a) MN-MYO; b) MYO; c) Sheets; d) No repair groups and are indicated by yellow stars. Scale bar – 100µm. e-e′) Representative image of a higher magnification view of mature NMJs (indicated by yellow stars) comprising of pretzel shaped AchR clusters (Purple) colabelling with presynaptic marker Synaptophysin (Red) located on the periphery of muscle fibers (Phalloidin-488). Scale bar – 10µm. f) Percentage of AchR clusters near the injury area (5mm^2^) that were innervated (Synaptophysin^+^) was counted across MN-MYO (n=5), MYO (n=4), Sheets (n=3) and No Repair (n=5) groups to depict maintenance/formation of mature NMJs in the host muscle. Mean percentage of mature NMJ of each group were as follows: MN-MYO – 78.95; MYO – 52.9; Sheets – 38.77; No Repair – 20.2. For indicated comparisons the individual p-values were as follows: MN-MYO vs MYO – p=0.0168 (*); MN-MYO vs Sheets – p=0.0012 (**); MN-MYO vs No Repair – p < 0.0001 (****). Error bars represent standard error of mean.

## Discussion

Severe musculoskeletal trauma like VML is accompanied by progressive motor axotomy over several weeks, leading to denervation of the injured muscle thereby severely limiting functional recovery^36^. Hence, appropriate somato-motor innervations remain one of the biggest challenges in fabricating a fully functional muscle. Apart from augmenting re-innervation process, tissue engineering strategies need to provide accurate cellular alignment and enable bulk muscle replacement to compensate for loss of muscle volume following VML. Aligned nanofiber scaffolds are the preferred biomaterial for muscle reconstruction since they not only promote myofibers alignment but can also be stacked to provide bulk to the engineered tissue. Aligned nanofiber scaffolds have been shown to facilitate NMJ formation *in vitro* as well as promote alignment of regenerating myofibers in vivo as compared to randomly oriented nanofibers^19,27,37^.

Most aligned nanofiber scaffolds used to date for VML repair are comprised of decellularized ECM or collagen which are prone to faster degradation and do not possess optimal mechanical properties to support organized myofibril regeneration. Additionally, synthetic polymer derived aligned nanofiber scaffolds are usually electropsun from polymer solutions thereby enabling scale-up and fabrication of custom designed sheets to fit the exact dimensions of an injured muscle. Although synthetic polymer derived aligned nanofibrous scaffolds have been shown to promote formation of functional NMJs in vitro^37^, they are yet to be used as scaffolds for VML repair. The present study is the first report on using synthetic polymer based aligned nanofiber scaffolds in a rat VML model. We have used commercially available nanofiber sheets made of polycaprolactone (PCL)-which is an FDA approved slowly degrading, bioresorbable polymer^38^. These aligned PCL nanofiber sheets were used as scaffolds for 3D motor neuron-myocyte coculture. We studied the effect of motor neurons on myocytes *in vitro* and observed that motor neurons cultured on pre-differentiated skeletal myocytes led to formation of mature NMJs and promoted fusion and bundling of myocytes to form multinucleate myofibers (Fig 3). This is in agreement with previous report which describes that enhanced fusion and maturation of myocytes are only observed when the myocytes are allowed to fully differentiate before introduction of the motor neurons and the coculture is maintained subsequently in serum-free conditions^5^.

To evaluate the *in vivo* potential of pre-innervated constructs as a reconstructive approach to VML, we used a standardized model of VML in the rat TA muscle^39^. Although different muscles like abdominal wall^40^, latissimus dorsi^41^ and quadriceps femoris^42,43^ have been used to create a VML grade critical muscle defect, the TA muscle remains the preferred choice of researchers due to ease of surgical access and measurable functional deficit following VML. Most tissue engineering strategies towards VML repair are evaluated in small animal models^44^. Cell based approaches generally comprise of cell lines (mouse/human) or primary cells and hence are carried out in athymic animals to allow in vivo survival and maturation of the constructs^19,27,44,45^. VML has been reported to lead to over 73% motor axotomy within 7 days post injury without significant change in the number of damaged axons up to 21 days^36^. This indicates that 7 days post injury is an appropriate acute time point to evaluate survival of implanted cells as well as study the effect of pre-innervated implants on host neuromuscular anatomy. We used T-cell deficient athymic rats to prevent immunogenic reaction to implanted mouse C2C12 cells and primary rat motor neurons. At terminal time point of 7 days post implant, the nanofiber sheets were still visible upon exposure of the TA and there were no apparent signs of immune rejection of the nanofiber sheets (Fig 4). The aligned nanofiber sheets were highly porous (80% porosity) and our method of stacking three layers of sheets allowed exchange of nutrients and oxygen through blood perfusion thereby facilitating survival of the implanted motor neurons and myocytes. Subsequent immunohistological analysis of transverse and longitudinal sections of the muscle allowed visualization of multiple layers nanofiber sheets and confirmed the presence of long thick bundles of skeletal myocytes and motor axons on the nanofiber sheets confirming acute survival of the implanted cells (Fig 5-6). In order to achieve a comprehensive understanding of the acute effects of innervation on the regenerative milieu of an injured muscle, we proceeded to investigate the density of satellite cells, microvasculature, AchR clusters and mature NMJs near the injury area.

The robust regenerative capacity of skeletal muscles can be largely attributed to the resident myogenic precursor cells called muscle satellite cells^30^. These satellite cells lie quiescent in between the basal lamina and sarcolemma and gets activated within a few days after an injury. Activated satellite cells then differentiate to form myoblasts which fuse together to form new skeletal muscle fiber^30^. Satellite cells can be reliably identified by paired box transcription factor Pax-7 which is expressed in both quiescent and activated stages^46^. Although pre-vascularized tissue engineered constructs have been shown to promote satellite cell activation upon implantation in a mild muscle injury model, the effects of pre-innervation on host satellite cell population are yet to be addressed^47^. Separate studies indicate that various neurotrophic factors like NGF and BDNF play a critical role in modulating satellite cell response within an injured muscle^48,49^. For instance, exogenous treatment with BDNF was enough to recover the regenerative capacity of satellite cells in BDNF-deficient mice after skeletal muscle injury^48^. Spinal motor neurons used in the present study for fabrication of pre-innervated constructs have been shown to secrete BDNF^50^. This can potentially explain the presence of significantly more Pax-7+ satellite cells near the injury area in MN-MYO group having pre-innrevated constructs comprising of motor neurons and myocytes (Fig 7). However, unlike Czajka *et al*’s report showing satellite cell migration within a pre-vascularized tissue engineered construct within 3 days of implantation, we did not observe any Pax-7+ satellite cells within our constructs by 7 days^47^ (Fig 7). This is likely due to the difference in models; VML presents a very different pathophysiology than does a mild incision injury. It is also possible that the inherent hydrophobic nature of the PCL nanofibers used here could have restricted host satellite cell infiltration^51,52^.

Tissue engineering strategies for VML repair demands bulk muscle reconstruction. Inadequate re-vascularization remains one of the major challenges to engineer thick skeletal muscle limiting nutrient exchange and survival of implanted cells^23^. Pre-vascularized constructs comprising of preformed vascular networks have been shown to promote microvasculature, vascular perfusion of the graft and inosculation with host vascular system thereby significantly improving muscle regeneration following VML^23,27,47^. Neurotrophic factors like NGF, BDNF, GDNF, NT-3 have been reported to enhance angiogenesis in different tissues like skin, heart and cartilage through receptor mediated activation or recruitment of proangiogenic precursor cells^53–55^. Spinal motor neurons secrete BDNF whereas astrocytes can express a range of neurotrophic factors^50,56^. It is to be noted that although we strive to obtain a pure motor neuron population, we have detected minimal glial cells in our motor neuron cultures (data not shown). We have observed that the pre-innervated constructs used in this study lead to significant increase in microvasculature near the injury area following implantation in a VML model (Fig 8). Although the molecular mechanisms of how pre-innervated constructs promote vascularization is the scope of future studies, it is reasonable to postulate, that neurotrophic factors from our spinal cord derived cell population (comprising of motor neurons and glia) triggered this increased microvasculature. Interestingly, despite such increased microvasculature near the injury area we did not find any evidence of endothelial cells within the implanted nanofiber sheets. Aside from the inherent hydrophobicity of PCL, the absence of endothelial cell infiltration within the graft can also indicate that the evaluation time of 7 days was too early to observe vascular integration of the construct. Muscle nicotinic AchRs are pentameric structures that are dispersed along the basal membrane of myofibers (extra-junctional) during fetal stage and progressively redistribute to form localized (junctional) pretzel shaped clusters on adult muscle^57^. AchR clustering plays a pivotal role in skeletal muscle function and regeneration through formation of stable motor end plates thereby effecting functional restoration following severe musculoskeletal trauma. Early physical rehabilitation involving exercise has been shown to benefit patients with VML in recovering muscle force and range of motions^58^. Similarly, rehabilitative exercise in conjunction with bioengineered constructs augments functional restoration in murine models of VML by promoting formation of AchR clusters and mature NMJs^59–62^. One of the key findings of the present study is that pre-innervated constructs augments AchR clustering and NMJ formation. This is reflected in our results which show MN-MYO group have significantly more AchR receptor clusters and mature NMJs within 7 days of implanting in a VML model (Fig 9-10). Clustering of AchRs in skeletal muscles is mainly controlled by motor innervation through secretion of neural agrin by the motor neurons^33,63^. This may be a reason behind increased density of AchR clusters near the injury area following implantation of pre-innervated constructs. In a denervated muscle following injury, AchR clusters can again disperse to form extra-junctional immature receptors. This often leads to an increase in overall count of Bungarotoxin+ structures in an injured denervated muscle^64^. Hence, in order to investigate if the observed increase in AchR clusters was only an injury effect, we looked for innervated AchR clusters that would indicate formation of mature NMJs and preservation of the motor end plate. The MN-MYO group was found to have significantly higher percentage of innervated AchR clusters as compared to other groups (Fig 10). This confirms that pre-innervated constructs promote formation of mature NMJs around the injury area at acute time points which can potentially have a significant effect in augmenting functional restoration at more chronic time points.

Although our study demonstrates the potential of pre-innervated constructs in promoting a regenerative environment following VML, several limitations exist. First, this study was conducted in athymic rats lacking an intact immune system and was terminated at an acute time point. Second, we speculate about the possible role of various neurotrophic factors in facilitating a pro-regenerative microenvironment. However, it’s very likely that the physical presence of preformed motor axonal network plays a crucial role. Hence, more in depth studies are necessary to elucidate the cellular/molecular mechanisms behind the observed effects of pre-innervation in VML repair. Third, although the benefits of preformed axonal networks for in vitro tissue engineering and acute host responses are apparent in our results, longer term effects of extraneous neurons on neuromuscular integration and functional recovery are unclear. To address these shortcomings, ongoing studies in our lab are looking at effects of pre-innervated constructs on functional recovery at more chronic time points.

In summary, the present study is the first to explore the implications of pre-innervation on the regenerative microenvironment in a rat VML model at an acute time point. This is also the first report on the use of synthetic polymer derived aligned nanofiber scaffolds as an implant in a VML model. Our results indicate that pre-innervation promotes myocyte maturation *in vitro*, satellite cell migration and vascularization in the injury area as well as facilitates formation of mature NMJs thereby providing a favorable regenerative microenvironment for neuromuscular regeneration following VML. We believe that these findings in skeletal muscle injury model would stimulate further research into developing pre-innervated tissue engineered constructs for application in smooth muscle as well as cardiac tissue engineering. In future work, these nerve-muscle constructs may also be fabricated using cells derived from adult human stem cell sources (e.g., iPSCs), thereby making them translational as an autologous, personalized bioengineered construct. These pro-regenerative effects can potentially lead to enhanced functional neuromuscular regeneration following VML, thereby improving the levels of functional recovery following these devastating injuries.

## Methods

### Isolation and Culture of Rat Spinal Motor Neurons

Motor neurons were harvested from the spinal cord of E16 Sprague Dawley rat embryos following previously described procedure^65,66^. All harvest procedures prior to dissociation were conducted on ice. Briefly, spinal cords were extracted from the pups and digested with 2.5% 10X trypsin diluted in 1mL L-15 for 15 mins at 37°C. The digested tissue was triturated multiple times with DNAse (1mg/mL) and 4% BSA and centrifuged at 280g for 10minutes to pool all the cell suspension. Subsequently, the cell suspension was subjected to Optiprep mediated density gradient centrifugation at 520g for 15 minutes to separate the motor neuron population. Following centrifugation, the supernatant was discarded, and cells were resuspended in motor neuron plating media consisting of glial conditioned media. Glial conditioned media was made as described earlier^65^ and supplemented with 37ng/mL hydrocortisone, 2.2 µg/mL isobutylmethylxanthine, 10 ng/mL BDNF, 10 ng/mL CNTF, 10 ng/mL CT-1, 10 ng/mL GDNF, 2% B-27, 20ng/mL NGF, 20 µM mitotic inhibitors, 2 mM L-glutamine, 417 ng/mL forskolin, 1 mM sodium pyruvate, 0.1 mM β-mercaptoethanol, 2.5 g/L glucose to make complete motor neuron plating media.

### Mouse Skeletal Myoblast Cell Line (C2C12) Culture

C2C12 cell line was maintained in growth media comprising of DMEM-High Glucose, supplemented with 20%FBS and 1% PennStrep. The cells were allowed to reach 80% confluency before inducing differentiation through differentiation media comprising of DMEM-High Glucose supplemented with 2% NHS and 1% Penicillin-Streptomycin.

### Motor Neuron-Myocyte Coculture on Nanofiber Sheets to form Pre-Innervated Tissue Engineered Muscle

A 15cm × 15cm PCL aligned nanofiber sheet was custom fabricated and purchased from Nanofiber Solutions LLC (Ohio, USA). The sheets were cut into 10mmx5mm pieces, placed in 24 well tissue culture plates and UV sterilized prior to coating with 20 µg/mL poly-D-lysine (PDL) in sterile cell culture water overnight. The sheets were subsequently washed thrice with PBS before coating with laminin (20 µg/mL) for 2 hours. Pre-differentiated C2C12 cells were plated on the nanofiber sheets at a concentration of 2×10^5^ cells/sheet in growth media for 24 hours before being cultured using differentiation media for 7 days in vitro (DIV) with regular changes of media. Dissociated motor neurons were plated on top of the myocyte layer at a concentration of 1×10^5^cells/sheet and the coculture was maintained with serum-free motor neuron media up to 14DIV with regular changes of media. The sheets with only myocytes were also kept on serum-free motor neuron media between 7-14DIV to maintain parity of cell culture condition between groups.

### Immunofluorescence staining of cell laden nanofiber sheets

Samples were fixed for 35 min in 4% paraformaldehyde (EMS, Cat# 15710), washed three times with 1× PBS, and permeabilized in 0.3% Triton-X100 + 4% Normal Horse Serum (NHS) (Sigma) for 60 min. Samples were blocked in 4% NHS (Sigma) and all subsequent steps were performed using 4% NHS for antibody dilutions. For staining of actin and AchR, samples were incubated with Alexfluor-488-conjugated phalloidin (1:200, Invitrogen, A12379) and AlexaFluor-647-conjugated bungarotoxin (1:250, Invitrogen, B35450). For assessment of motor neuron morphology and maturity, separate fixed samples were incubated with an axonal marker Tuj-1 (1:250, Abcam, ab18207) and presynaptic marker Synaptophysin (1:500, abcam, ab32127) for 16 h at 4 °C followed by Alexa Fluor-568 antibody (Life Technologies). Images were acquired using a Nikon Eclipse TI A1RSI laser scanning confocal microscope.

### Quantification of Myocyte Fusion Index

Multiple replicates of MN-MYO (n=7) and MYO only (n=14) cultures were considered for measuring myocyte fusion index (MFI) as per the following equation -

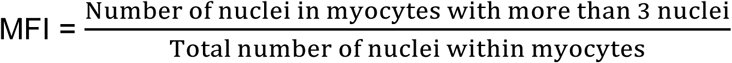

At least three 2mm^2^ area was considered per sample for counting MFI and the average was plotted for each sample (Fig 3c).

### Bioscaffold Implantation in Rat Model of VML

All procedures were approved by the Institutional Animal Care and Use Committee at the Michael J. Crescenz VA Medical Center and adhered to guidelines established in the PHS Policy on Humane Care and Use of Laboratory Animals (2015). Rats had access to food and water ad libitum and were pair-housed in a colony with a 12hr light/dark cycle.

Adult male athymic rats (RNU strain 316; Charles River Labs) weighing 280-300g were used as subjects for this study. All procedures were carried out under aseptic conditions while the animal was under general anesthesia (1.5-2% isoflurane, 1.5L O2) and thermal support was provided via temperature-controlled water pad. After shaving the hair of the lower left hind limb and applying a liberal coat of betadine solution, 0.25 mg of bupivacaine was administered subcutaneously along the planned incision line. Following a previously outlined procedure^39^, a longitudinal skin incision was made along the lateral aspect of the lower leg; care was taken not to cut through the underlying fascia covering the tibialis anterior (TA) muscle. The skin was bluntly dissected from the fascia along the length of the TA. A longitudinal incision (~1.5cm) was made in the overlying fascia and the fascia was then gently dissected from the underlying TA muscle using a blunt probe, keeping the fascia intact for later repair. Once the muscle was exposed, a flat spatula was inserted between the tibial bone and TA muscle in order to isolate the TA/extensor digitorum longus (EDL) complex for further surgical manipulation. A mark was made 0.5cm from the tibial tuberosity, indicating the proximal incision in the TA. A second mark was made 1.0cm distal to the first and a 1.0cm X 0.7cm area was outlined on the TA using a surgical caliper (Fine Science Tools, cat # 18000-35). A 3cm deep incision was made through the muscle at the proximal line and the scalpel turned parallel to the tibial bone to make a smooth cut through the muscle while following the outlined rectangle. Care was taken to avoid cutting completely through the TA or slicing the underlying EDL muscle. Once the portion of the muscle was removed, it was weighed and discarded. Deficits were repaired with 3 stacked cell laden sheets (MN-MYO, MYO), 3 stacked acellular sheets alone (SHEETS), or were not repaired (NO REPAIR). Prior to implantation, the sheets were washed thoroughly with PBS to remove any leftover media. The fascia, connective tissue, and skin were closed in layers with 8-0 prolene, 6-0 prolene, or staples, respectively. At the conclusion of the surgery, the area was cleaned with alcohol and animals were given a subcutaneous injection of sustained-release meloxicam (4mg/kg). Animals were placed on heating pads until recovered and returned to home cages.

### Immunohistological Assessment of Injury Site at Acute Time Point

Freshly harvested whole anterior muscle samples (TA+EDL) were fixed in 4% paraformaldehyde (EMS, Cat# 15710), submerged in 20% sucrose in 1X phosphate buffered saline (PBS, pH 7.4) for density equilibration, frozen and cryosectioned axially (20um) across the middle portion of the graft region. Prior to staining, sections were washed three times in 1X PBS, blocked and permeablized in 4% normal horse serum (Sigma, G6767) with 0.3% Triton X-100 (Sigma, T8787) in 1X PBS for one hour. All subsequent steps were performed using blocking solution for antibody dilutions. For staining of skeletal muscle actin, samples were incubated with rabbit-anti-skeletal muscle actin (1:500, abcam, ab46805) overnight at 4°C followed by AlexaFluor-568 antibody (1:500, Invitrogen, A10042) for two hours at room temperature. Alternatively, for staining of actin, samples were incubated with AlexaFluor-488-conjugated phalloidin (1:400, Invitrogen, A12379) for two hours at room temperature. For microvasculature staining, smooth muscle actin and endothelial cells were targeted, samples were incubated with mouse-anti-smooth muscle actin (1:500, abcam, ab7817) or rabbit-anti-CD31/PECAM1 (1:500, Novus, NB100-2284) overnight at 4°C followed by AlexaFluor-568 antibody (1:500) or AlexaFluor-568 antibody (1:500, Invitrogen, A10087), respectively, for two hours at room temperature. For staining of satellite cells, samples were incubated with mouse-anti-Pax7 (1:10, DSHB) overnight at 4°C followed by AlexaFluor-647 antibody (1:500, Invitrogen, A31573) for two hours at room temperature. For staining of axons, samples were incubated with sheep-anti-ChAT (1:500, abcam, ab18736) or rabbit-anti-NF200 (1:500, abcam, ab8135) overnight at 4°C followed by AlexaFluor-568 (1:500, Invitrogen, A21099 and AlexaFluor-647 antibody (1:500, Invitrogen, A31573) respectively, for two hours at room temperature. For staining of laminin, samples were incubated with rabbit-anti-laminin (1:500, abcam, ab11575) overnight at 4°C followed by AlexaFluor-568 antibody (1:500) for two hours at room temperature. For staining of neuromuscular junctions, samples were co-labeled with synaptophysin and bungarotoxin. Samples were incubated overnight with rabbit-anti-synaptophyhsin (1:500, abcam, ab32127) at 4°C followed by AlexaFluor-568 antibody (1:500) and concurrently with AlexaFluor-647-conjugated bungarotoxin (1:1000, Invitrogen, B35450) for two hours at room temperature. For staining of cell nuclei, samples were incubated with Hoescht (1:10,000, Invitrogen, H3570) for 20 minutes at room temperature. Images were acquired using a Nikon Eclipse TI A1RSI laser scanning confocal microscope.

For quantitative measurement of satellite cell, micro-vessel, AchR cluster and mature NMJ density, an area of 5mm^2^ (5mm long and 1mm wide) was chosen at 100µm from injury/implant site towards the host muscle and defined as the injury area. At least 3 cross-sections each separated by 300µm was considered for counting and average density was plotted in the graph and compared across groups.

### Statistical Analysis

All quantifications reported in this study were performed by personnel blinded about the treatment groups. All statistical analysis was performed using GraphPad PRISM software. For comparison between two groups only (Fig 3c), an unpaired two-tailed Student’s t-test with Welch’s correction was used. For comparison between multiple groups, a one-way analysis of variance (ANOVA) was performed with post hoc Tukey’s adjustment with 95% Confidence Interval (Fig 7f, 8f, 9f, 10f). Significance was taken at p ≤ 0.05 (*), p ≤ 0.01 (**), p ≤ 0.001 (***), and p ≤ 0.0001 (****). All graphs were made in GraphPad PRISM and display mean ± standard error of mean (SEM).

## Data Availability

Data supporting the conclusions of this paper are available from the corresponding author upon reasonable request.

## Author Contributions

S.D., K.D.B., F.A.L., and D.K.C. designed and carried out experiments and analyzed data. S.D, K.D.B and D.K.C interpreted the results. F.A.L, J.C.M. and H.K. performed histological experiments and assisted with blinded quantification. K.D.B. performed the surgeries, and F.M. and C.A. provided guidance. S.D. and D.K.C. wrote and organized the manuscript, with editorial input from K.D.B, F.A.L., F.M., C.A. and Z.S.A. D.K.C. conceived of the approach.

### Acknowledgements

Financial support was provided by the U.S. Department of Defense through the Medical Research and Materiel Command [W81XWH-15-1-0466 (Cullen) & W81XWH-16-1-0796 (Cullen)], the National Institutes of Health [BRAIN Initiative U01-NS094340 (Cullen)], and the Department of Veterans Affairs [Merit Review I01-BX003748 (Cullen)]. Any opinion, findings, and conclusions or recommendations expressed in this material are those of the authors(s) and do not necessarily reflect the views of the Department of Defense, National Institutes of Health, or Department of Veterans Affairs.

## References

1. Kreipke, R. E. & Birren, S. J. Innervating sympathetic neurons regulate heart size and the timing of cardiomyocyte cell cycle withdrawal. J. Physiol. (2015). doi:10.1113/JP270917

2. Andreassen, A. K. Point:Counterpoint: Cardiac denervation does/does not play a major role in exercise limitation after heart transplantation. J. Appl. Physiol..104, 559–560 (2008).

3. Oberpenning, F., Meng, J., Yoo, J. J. & Atala, A. De novo reconstitution of a functional mammalian urinary bladder by tissue engineering. Nat. Biotechnol. (1999). doi:10.1038/6146

4. Morimoto, Y., Kato-Negishi, M., Onoe, H. & Takeuchi, S. Three-dimensional neuron-muscle constructs with neuromuscular junctions. Biomaterials (2013). doi:10.1016/j.biomaterials.2013.08.062

5. Guo, X., Gonzalez, M., Stancescu, M., Vandenburgh, H. H. & Hickman, J. J. Neuromuscular junction formation between human stem cell-derived motoneurons and human skeletal muscle in a defined system. Biomaterials (2011). doi:10.1016/j.biomaterials.2011.09.014

6. Das, M., Rumsey, J. W., Bhargava, N., Stancescu, M. & Hickman, J. J. A defined long-term in vitro tissue engineered model of neuromuscular junctions. Biomaterials (2010). doi:10.1016/j.biomaterials.2010.02.055

7. Uzel, S. G. M. et al. Microfluidic device for the formation of optically excitable, three-dimensional, compartmentalized motor units. Sci. Adv 2, (2016).

8. Wu, X., Corona, B. T., Chen, X. & Walters, T. J. A Standardized Rat Model of Volumetric Muscle Loss Injury for the Development of Tissue Engineering Therapies. Biores. Open Access (2012). doi:10.1089/biores.2012.0271

9. Turner, N. J. & Badylak, S. F. Regeneration of skeletal muscle. Cell and Tissue Research (2012). doi:10.1007/s00441-011-1185-7

10. Larouche, J., Greising, S. M., Corona, B. T. & Aguilar, C. A. Robust inflammatory and fibrotic signaling following volumetric muscle loss: A barrier to muscle regeneration comment. Cell Death and Disease (2018). doi:10.1038/s41419-018-0455-7

11. Grogan, B. F. & Hsu, J. R. Volumetrie muscle loss. J. Am. Acad. Orthop. Surg. (2011). doi:10.5435/00124635-201102001-00007

12. Corona, B. T., Rivera, J. C., Owens, J. G., Wenke, J. C. & Rathbone, C. R. Volumetric muscle loss leads to permanent disability following extremity trauma. J. Rehabil. Res. Dev. (2015). doi:10.1682/JRRD.2014.07.0165

13. Chuang, D. C. C. Free tissue transfer for the treatment of facial paralysis. Facial Plast. Surg. (2008). doi:10.1055/s-2008-1075834

14. Mertens, J. P., Sugg, K. B., Lee, J. D. & Larkin, L. M. Engineering muscle constructs for the creation of functional engineered musculoskeletal tissue. Regen. Med. (2014). doi:10.2217/rme.13.81

15. Zhang, J. et al. Perfusion-decellularized skeletal muscle as a three-dimensional scaffold with a vascular network template. Biomaterials (2016). doi:10.1016/j.biomaterials.2016.02.040

16. Sicari, B. M. et al. An acellular biologic scaffold promotes skeletal muscle formation in mice and humans with volumetric muscle loss. Sci. Transl. Med. (2014). doi:10.1126/scitranslmed.3008085

17. Grasman, J. M., Zayas, M. J., Page, R. L. & Pins, G. D. Biomimetic scaffolds for regeneration of volumetric muscle loss in skeletal muscle injuries. Acta Biomaterialia (2015). doi:10.1016/j.actbio.2015.07.038

18. Aurora, A., Roe, J. L., Corona, B. T. & Walters, T. J. An acellular biologic scaffold does not regenerate appreciable de novo muscle tissue in rat models of volumetric muscle loss injury. Biomaterials (2015). doi:10.1016/j.biomaterials.2015.07.040

19. Nakayama, K. H. et al. Rehabilitative exercise and spatially patterned nanofibrillar scaffolds enhance vascularization and innervation following volumetric muscle loss. npj Regen. Med. (2018). doi:10.1038/s41536-018-0054-3

20. Greising, S. M. et al. Unwavering Pathobiology of Volumetric Muscle Loss Injury. Sci. Rep. (2017). doi:10.1038/s41598-017-13306-2

21. Gilbert-Honick, J. et al. Engineering functional and histological regeneration of vascularized skeletal muscle. Biomaterials (2018). doi:10.1016/j.biomaterials.2018.02.006

22. Corona, B. T., Wenke, J. C. & Ward, C. L. Pathophysiology of volumetric muscle loss injury. Cells Tissues Organs (2016). doi:10.1159/000443925

23. Levenberg, S. et al. Engineering vascularized skeletal muscle tissue. Nat. Biotechnol. (2005). doi:10.1038/nbt1109

24. Aguilar, C. A. et al. Multiscale analysis of a regenerative therapy for treatment of volumetric muscle loss injury. Cell Death Discov. (2018). doi:10.1038/s41420-018-0027-8

25. Corona, B. T., Rivera, J. C., Wenke, J. C. & Greising, S. M. Tacrolimus as an adjunct to autologous minced muscle grafts for the repair of a volumetric muscle loss injury. J. Exp. Orthop. (2017). doi:10.1186/s40634-017-0112-6

26. Garg, K., Corona, B. T. & Walters, T. J. Losartan administration reduces fibrosis but hinders functional recovery after volumetric muscle loss injury. J. Appl. Physiol. (2014). doi:10.1152/japplphysiol.00689.2014

27. Nakayama, K. H. et al. Treatment of volumetric muscle loss in mice using nanofibrillar scaffolds enhances vascular organization and integration. Commun. Biol. (2019). doi:10.1038/s42003-019-0416-4

28. Quarta, M. Volumetric muscle loss: Including nerves into the equation. Muscle and Nerve (2018). doi:10.1002/mus.26080

29. Ferraro, E., Molinari, F. & Berghella, L. Molecular control of neuromuscular junction development. Journal of Cachexia, Sarcopenia and Muscle (2012). doi:10.1007/s13539-011-0041-7

30. Relaix, F. & Zammit, P. S. Satellite cells are essential for skeletal muscle regeneration: the cell on the edge returns centre stage. Development (2012). doi:10.1242/dev.069088

31. Yin, H., Price, F. & Rudnicki, M. A. Satellite cells and the muscle stem cell niche. Physiol. Rev. (2013). doi:10.1152/physrev.00043.2011

32. Wu, H., Xiong, W. C. & Mei, L. To build a synapse: signaling pathways in neuromuscular junction assembly. Development (2010). doi:10.1242/dev.038711

33. Scott, J. B. et al. Achieving Acetylcholine Receptor Clustering in Tissue-Engineered Skeletal Muscle Constructs In vitro through a Materials-Directed Agrin Delivery Approach. Front. Pharmacol. 7, 508 (2017).

34. Blumenthal, E. M., Conroy, W. G., Romano, S. J., Kassner, P. D. & Berg, D. K. Detection of functional nicotinic receptors blocked by α-bungarotoxin on PC12 cells and dependence of their expression on post-translational events. J. Neurosci. (1997).

35. Boido, M. & Vercelli, A. Neuromuscular Junctions as Key Contributors and Therapeutic Targets in Spinal Muscular Atrophy. Front. Neuroanat. (2016). doi:10.3389/fnana.2016.00006

36. Corona, B. T. et al. Impact of volumetric muscle loss injury on persistent motoneuron axotomy. Muscle and Nerve (2018). doi:10.1002/mus.26016

37. Luo, B. et al. Electrospun nanofibers facilitate better alignment, differentiation, and long-term culture in an: In vitro model of the neuromuscular junction (NMJ). Biomater. Sci. (2018). doi:10.1039/c8bm00720a

38. Guarino, V., Gentile, G., Sorrentino, L. & Ambrosio, L. Polycaprolactone: Synthesis, Properties, and Applications. in. Encyclopedia of Polymer Science and Technology 1–36 (John Wiley & Sons, Inc., 2017). doi:10.1002/0471440264.pst658

39. Wu, X., Corona, B. T., Chen, X. & Walters, T. J. A Standardized Rat Model of Volumetric Muscle Loss Injury for the Development of Tissue Engineering Therapies. Biores. Open Access 1, 280–290 (2012).

40. Conconi, M. T. et al. Homologous muscle acellular matrix seeded with autologous myoblasts as a tissue-engineering approach to abdominal wall-defect repair. Biomaterials (2005). doi:10.1016/j.biomaterials.2004.07.035

41. Passipieri, J. A. et al. In Silico and In Vivo Studies Detect Functional Repair Mechanisms in a Volumetric Muscle Loss Injury. Tissue Eng. Part A (2019). doi:10.1089/ten.tea.2018.0280

42. Li, M. T. A., Willett, N. J., Uhrig, B. A., Guldberg, R. E. & Warren, G. L. Functional analysis of limb recovery following autograft treatment of volumetric muscle loss in the quadriceps femoris. J. Biomech. (2014). doi:10.1016/j.jbiomech.2013.10.057

43. Mase, V. J. et al. Clinical Application of an Acellular Biologic Scaffold for Surgical Repair of a Large, Traumatic Quadriceps Femoris Muscle Defect. Orthopedics (2010). doi:10.3928/01477447-20100526-24

44. Vigodarzere, G. C. & Mantero, S. Skeletal muscle tissue engineering: Strategies for volume tric constructs. Frontiers in Physiology (2014). doi:10.3389/fphys.2014.00362

45. Koffler, J. et al. Improved vascular organization enhances functional integration of engineered skeletal muscle grafts. Proc. Natl. Acad. Sci. (2011). doi:10.1073/pnas.1017825108

46. Sambasivan, R. et al. Pax7-expressing satellite cells are indispensable for adult skeletal muscle regeneration. Development (2011). doi:10.1242/dev.073601

47. Czajka, C. A., Calder, B. W., Yost, M. J. & Drake, C. J. Implanted scaffold-free prevascularized constructs promote tissue repair. Ann. Plast. Surg. (2015). doi:10.1097/SAP.0000000000000439

48. Clow, C. & Jasmin, B. J. Brain-derived Neurotrophic Factor Regulates Satellite Cell Differentiation and Skeltal Muscle Regeneration. Mol. Biol. Cell (2010). doi:10.1091/mbc.e10-02-0154

49. Lavasani, M., Lu, A., Peng, H., Cummins, J. & Huard, J. Nerve Growth Factor Improves the Muscle Regeneration Capacity of Muscle Stem Cells in Dystrophic Muscle. Hum. Gene Ther. (2006). doi:10.1089/hum.2006.17.ft-175

50. Joseph, M. S., Tillakaratne, N. J. K. & de Leon, R. D. Treadmill training stimulates brain-derived neurotrophic factor mRNA expression in motor neurons of the lumbar spinal cord in spinally transected rats. Neuroscience (2012). doi:10.1016/j.neuroscience.2012.08.024

51. Tsai, W. B., Chen, C. H., Chen, J. F. & Chang, K. Y. The effects of types of degradable polymers on porcine chondrocyte adhesion, proliferation and gene expression. J. Mater. Sci. Mater. Med. (2006). doi:10.1007/s10856-006-8234-x

52. Ishaug-Riley, S. L., Okun, L. E., Prado, G., Applegate, M. A. & Ratcliffe, A. Human articular chondrocyte adhesion and proliferation on synthetic biodegradable polymer films. Biomaterials (1999). doi:10.1016/S0142-9612(99)00155-6

53. Kermani, P. & Hempstead, B. BDNF: A Newly Described Mediator of Angiogenesis. Trends Cardiovasc Med. (2008). doi:10.1016/j.tcm.2007.03.002

54. Blais, M., Lévesque, P., Bellenfant, S. & Berthod, F. Nerve Growth Factor, Brain-Derived Neurotrophic Factor, Neurotrophin-3 and Glial-Derived Neurotrophic Factor Enhance Angiogenesis in a Tissue-Engineered In Vitro Model. Tissue Eng. Part A (2013). doi:10.1089/ten.tea.2012.0745

55. Lin, C. Y. et al. Brain-derived neurotrophic factor increases vascular endothelial growth factor expression and enhances angiogenesis in human chondrosarcoma cells. Biochem. Pharmacol. (2014). doi:10.1016/j.bcp.2014.08.008

56. Toyomoto, M. et al. Production of NGF, BDNF and GDNF in mouse astrocyte cultures is strongly enhanced by a cerebral vasodilator, ifenprodil. Neurosci. Lett. (2005). doi:10.1016/j.neulet.2004.12.063

57. Fuhrer, C. & Huh, K.-H. Clustering of Nicotinic Acetylcholine Receptors: From the Neuromuscular Junction to Interneuronal Synapses. Mol. Neurobiol. (2003). doi:10.1385/mn:25:1:079

58. Dziki, J. et al. An acellular biologic scaffold treatment for volumetric muscle loss: results of a 13-patient cohort study. npj Regen. Med. (2016). doi:10.1038/npjregenmed.2016.8

59. Quarta, M. et al. Bioengineered constructs combined with exercise enhance stem cell-mediated treatment of volumetric muscle loss. Nat. Commun. (2017). doi:10.1038/ncomms15613

60. Greising, S. M. et al. Early rehabilitation for volumetric muscle loss injury augments endogenous regenerative aspects of muscle strength and oxidative capacity. BMC Musculoskelet. Disord. (2018). doi:10.1186/s12891-018-2095-6

61. Aurora, A., Garg, K., Corona, B. T. & Walters, T. J. Physical rehabilitation improves muscle function following volumetric muscle loss injury. BMC Sports Sci. Med. Rehabil. (2014). doi:10.1186/2052-1847-6-41

62. Gentile, N. E. et al. Targeted Rehabilitation after extracellular matrix scaffold transplantation for the treatment of volumetric muscle loss. Am. J. Phys. Med. Rehabil. (2014). doi:10.1097/PHM.0000000000000145

63. Reist, N. E., Werle, M. J. & McMahan, U. J. Agrin released by motor neurons induces the aggregation of acetylcholine receptors at neuromuscular junctions. Neuron (1992). doi:10.1016/0896-6273(92)90200-W

64. Edwards, C. The effects of innervation on the properties of acetylcholine receptors in muscle. Neuroscience 4, 565–584 (1979).

65. Katiyar, K. S., Struzyna, L. A., Das, S. & Cullen, D. K. “Stretch-Growth” of Motor Axons in Custom Mechanobioreactors to Generate Long-Projecting Axonal and Axonal-Myocyte Constructs. bioRxiv 598755 (2019). doi:10.1101/598755

66. Graber, D. J. & Harris, B. T. Purification and culture of spinal motor neurons from rat embryos. Cold Spring Harb. Protoc. (2013). doi:10.1101/pdb.prot074161

